# Semaglutide, Tirzepatide, and Retatrutide Attenuate the Interoceptive Effects of Alcohol in Male and Female Rats

**DOI:** 10.1101/2025.04.17.649402

**Authors:** McKinley Windram, Dennis F. Lovelock, Joseph M. Carew, Caroline G. Krieman, Christian S. Hendershot, Joyce Besheer

## Abstract

**Rationale:** Alcohol use disorder (AUD) remains a major public health challenge, yet effective pharmacotherapies are limited. As such, there is growing interest in repurposing medications with novel mechanisms of action. Glucagon-like peptide-1 (GLP-1) receptor agonists, originally developed for type 2 diabetes, have emerged as promising candidates due to effects on intake regulation and reward processing. GLP-1 receptor agonists, including semaglutide, reduce alcohol intake and relapse-like behaviors in rodent and non-human primate models, and a recent clinical trial found that semaglutide decreased alcohol craving and drinking in adults with AUD. Modulation of the subjective/interoceptive effects of alcohol may contribute to the therapeutic potential of GLP-1 receptor agonists.

**Objectives:** This study used operant drug discrimination in male and female rats to assess how acute and repeated semaglutide treatment affects alcohol’s discriminative stimulus (interoceptive) effects. We hypothesized that GLP-1 receptor activation would disrupt alcohol’s interoceptive effects. We also evaluated acute treatment with tirzepatide, a dual GLP-1/gastric inhibitory peptide (GIP) receptor agonist, and retatrutide, a triple GIP/GLP-1/glucagon receptor agonist, to determine whether broader receptor activity would differentially influence alcohol’s subjective effects.

**Results:** Acute administration of semaglutide, tirzepatide, and retatrutide each attenuated alcohol discrimination, suggesting modulation of subjective alcohol effects. Repeated semaglutide maintained efficacy across the 15-day treatment period; alcohol discrimination returned to control levels three days after treatment cessation.

**Conclusions:** Building on prior work with GLP-1 receptor agonists, these results provide important context for interpreting clinical observations of reduced drinking behavior among individuals receiving this class of therapeutics.

## Introduction

Alcohol use disorder (AUD) remains a major public health challenge, affecting over 10% of the U.S. population aged 12 and older (SAMSHA, 2023). Despite its prevalence, effective pharmacological treatments are limited. Current FDA-approved treatments, such as disulfiram, acamprosate and naltrexone, can help manage AUD by reducing consumption and relapse risk, but their effectiveness varies widely across individuals (Heilig et al. 2024). Given the heterogeneity of AUD and the complexity of the effect of alcohol on neural and physiological systems, there is a critical need to explore alternative pharmacotherapies that act through novel mechanisms. One promising avenue is the use of incretin medications, particularly glucagon-like peptide-1 (GLP-1) receptor agonists, which have shown potential in preclinical and clinical studies (Fink-Jensen and Vilsbøll 2016; Hayes and Schmidt 2016; Brunchmann et al. 2019; Klausen et al. 2022; Jerlhag 2023, 2025; Leggio et al. 2023; Bruns et al. 2024). Repurposing of long-acting FDA-approved GLP-1 receptor agonists such as semaglutide, the dual GLP-1/gastric inhibitory polypeptide (GIP) receptor agonist tirzepatide, and the triple-agonist retatrutide which additionally targets glucagon receptors and is currently in clinical trials for obesity, may offer a promising strategy treating AUD.

GLP-1 receptor agonists were initially developed to improve glycemic control in type 2 diabetes by enhancing insulin secretion and inhibiting glucagon release, with additional effects on appetite regulation and gastric emptying later recognized as clinically relevant (Chao et al. 2023; Badulescu et al. 2024). However, GLP-1 receptors are also expressed in brain regions involved in reward processing, and a GLP-1R variant was found to be associated with increased alcohol use, suggesting a potential role in modulating drug- and alcohol-seeking behavior (Merchenthaler et al. 1999; Hayes and Schmidt 2016; Tufvesson-Alm et al. 2023; Badulescu et al. 2024; Heilig et al. 2024; Kooij et al. 2024). Preclinical studies have demonstrated the efficacy of short-acting and the more recent long-acting GLP-1 receptor agonists to reduce alcohol self-administration, relapse-like behaviors, and binge drinking in rodent models (Egecioglu et al. 2013; Suchankova et al. 2015; Vallöf et al. 2016; Marty et al. 2020; Klausen et al. 2022; Aranäs et al. 2023, 2025; Chuong et al. 2023). Similarly, in non-human primates, GLP-1 receptor agonists including semaglutide have been shown to suppress alcohol intake (Suchankova et al. 2015; Thomsen et al. 2019; Fink-Jensen et al. 2024). Observational studies in humans being treated for other conditions have noted incidental reductions in alcohol use or related outcomes among patients receiving GLP-1 receptor agonists (Quddos et al. 2023; Richards et al. 2023; Wang et al. 2024), and the first randomized controlled trial of semaglutide in AUD found that low-dose treatment reduced alcohol craving and drinking quantity, providing evidence for its therapeutic potential (Hendershot et al. 2025).

The growing popularity of GLP-1 receptor agonists, especially semaglutide (Ozempic, Wegovy), has been accompanied by considerable social media attention and discussion, from which thorough analyses of user-generated data have emerged highlighting reductions in alcohol motivation, decreased drinking behavior, and altered subjective responses to alcohol (Arillotta et al. 2023, 2024; Quddos et al. 2023; Bremmer and Hendershot 2024). We will refer to these subjective responses as interoceptive effects of alcohol. These subjective/interoceptive effects are crucial because they have the potential to significantly influence drinking behavior, motivation to drink, and relapse risk (Solinas et al. 2006; Koob and Volkow 2010; Morean and Corbin 2010; Ray et al. 2010; King et al. 2014, 2016; Jaramillo et al. 2018b; Lovelock et al. 2021). If GLP-1 receptor activation disrupts or modifies the processing of these internal cues, it may represent a critical mechanism underlying the observed reductions in alcohol motivation and consumption. Drug discrimination procedures are well-established in both human and preclinical studies for assessing subjective/interoceptive effects (Grant and Bennett 2003; Bolin et al. 2016; Lovelock et al. 2021), making them particularly valuable for evaluating whether GLP-1 agonists influence the subjective perception of the pharmacological properties of alcohol. Investigating this mechanism may offer valuable insights into how GLP-1 receptor agonists modulate alcohol-related behaviors and their therapeutic utility in treating AUD.

To date, characterization of the effects of GLP-1 agonists on the subjective/interoceptive effects of alcohol has not been reported. In this paper, we used operant drug discrimination procedures in male and female rats where alcohol (2 g/kg) was trained as a discriminative stimulus. We hypothesized that semaglutide pretreatment would disrupt the discriminative stimulus effects of alcohol following acute and repeated treatment, indicating altered interoceptive effects of alcohol. We further examined whether acute tirzepatide, a dual glucose-dependent insulinotropic polypeptide (GIP) and GLP-1 receptor agonist recently approved by the FDA for obesity, and retatrutide, an investigational triple agonist targeting GIP, GLP-1, and glucagon receptors currently in Phase 3 trials would recapitulate or enhance these effects. Clarifying these interactions may inform the clinical use and further development of GLP-1 receptor agonists as effective treatments for AUD.

## Methods

### Subjects

Adult male and female Long–Evans rats (Envigo-Harlan, Indianapolis, IN) were delivered at 7 weeks old and housed in same-sex pairs in ventilated cages. Two separate cohorts of rats were used in these studies. Cohort 1 (n=7 males, 9 females) was used for Experiment 1 and cohort 2 (n=7 males, 3 females) was used in Experiments 2-4. The colony room was maintained on a 12-hour light/dark schedule, and all experiments were conducted during the light portion of the schedule. Rats were fed daily to maintain body weight at 90% of free-feeding weight and had ad libitum access to water. Animals were under continuous care and monitoring by veterinary staff from the Division of Comparative Medicine (DCM) at UNC-Chapel Hill. All procedures were carried out in accordance with the NIH Guide for Care and Use of Laboratory Animals and institutional guidelines. All protocols were approved by the UNC Institutional Animal Care and Use Committee (IACUC). UNC-Chapel Hill is accredited by the Association for Assessment and Accreditation of Laboratory Animal Care (AAALAC).

### Drugs

Alcohol solutions were prepared by diluting ethanol (95% v/v; Pharmco-AAPER, Shelbyville, KY) in distilled water to a 20% v/v concentration. Semaglutide, tirzepatide, and retatrutide (Biosynth, Compton, UK, lot #s 0000042482, 0000054911, and 0000205204 respectively) were dissolved in 0.3% phosphate-buffered saline (PBS), which was adjusted to a pH of 8.0 using NaOH and injected subcutaneously at 1 ml/kg.

### Apparatus

Standard rat operant chambers (Med Associates, Inc., Fairfax VT) measuring 31 × 24 × 32 cm were located inside noise dampening cubicles that contained fans which provided ventilation and masked sound. Each chamber was equipped with two levers positioned on either side of a liquid dipper mechanism, with cue lights installed above each lever. Completion of a fixed ratio 10 (FR10) schedule of reinforcement on the appropriate lever activated a dipper that presented 0.1 mL of sucrose (10% w/v) and illuminated the cue light above the lever for 4 seconds. Responses on the other lever were recorded but produced no programmed consequence.

### Lever training

Initially, rats underwent overnight lever training sessions as previously described (Besheer and Hodge 2005; Jaramillo et al. 2016, 2018a; Tyler et al. 2022, 2024). Briefly, in these 16-hour overnight sessions one lever (e.g., left or right) was available and a 10% sucrose reward was offered starting at a fixed ratio of 1 (FR1) for every lever press which escalated to FR4 by the end of the session. Rats experienced two of these sessions – one with each lever. Next, to train the rats to respond on each lever at FR10, they underwent five-days-a-week training sessions of 40 minutes each across five weeks. During these sessions, only one lever was available at a time, with the available lever alternating between the right and left every second session. Initially, sucrose was delivered at an FR4 schedule; following 4–8 training days, the response requirement was incrementally increased by two (to FR6, FR8, etc.), until 8 consecutive days of responding at FR10.

### Operant discrimination training

Next, discrimination training began in daily training sessions (M-F) where either 2 g/kg of a 20% alcohol solution (v/v) or a volume-equivalent amount of water was administered (i.g.). Immediately after the alcohol or water administration, rats were placed in the chambers and after a 20-minute delay, the house light was illuminated, and both levers were introduced into the chamber signaling the start of the 15-minute training session. After alcohol administration, 10 responses on the “alcohol-appropriate” lever (e.g., “correct” lever) resulted in sucrose delivery, while responses on the other (e.g., “incorrect” lever) produced no programmed consequence. After water administration, completion of 10 responses on the water-appropriate lever resulted in sucrose delivery. Lever contingencies were counterbalanced such that half of the rats were trained with the left lever as the alcohol-appropriate lever and the other half with the right lever. Alcohol and water sessions alternated every two days. Training sessions continued until the percent of alcohol-and water-appropriate lever responses before the first reinforcer and during the entire session was >80% for 8 of 10 consecutive. Once these criteria were met, testing of an alcohol dose response curve began.

### Discrimination testing

First, an alcohol dose response curve (0.5, 1.0, 2.0, 2.5 g/kg, i.g.) was generated to confirm discriminative stimulus control of behavior. Test sessions were identical to training sessions except that they were 2-min in duration and FR10 on either lever resulted in sucrose delivery to prevent lever selection bias and to allow for quantification of response rate. Once the alcohol dose response curve was completed, testing of the compounds began. Assigned drug doses or vehicle were administered 3 h prior to the test session for all compounds tested.

### Experiment 1: Semaglutide

Sixteen rats (7 males and 9 females) were initially assigned in a counterbalanced manner to one of four treatment conditions: 0, 0.01, 0.03, or 0.1 mg/kg semaglutide. Each rat underwent an 18-day cycle during which they received the assigned injection every other day—totaling 8 injections over 15 days—to simulate once-weekly administration in human patients (Geisler 2022). Test sessions were conducted on the first, third, and eighth injection days (termed injection 1, injection 3, and injection 8, respectively) and on post-treatment days 1 and 3 (see Figure 3A), and training sessions were withheld except for the test sessions. On the test sessions, semaglutide or vehicle was administered 3 h prior to alcohol (2 g/kg, IG), with the exception of the post-treatment tests in which only alcohol was administered. This cycle was repeated four times, ensuring that each rat was exposed to every dose, which allowed for within-subject comparisons. Between each testing cycle, rats completed a minimum of 10 training days to serve as a washout period and to reinforce accurate discrimination performance. Following the fourth cycle, a single test session was conducted using a dose of 0.003 mg/kg semaglutide to serve as a comparison with the initial test at each dose. Data were analyzed based on three criteria: acute administration (injection 1), repeated administration (injections 1, 3, and 8), and post-treatment effects (days 1 and 3 post-treatment).

### Experiment 2: Semaglutide administered on water days

To determine whether the effects of semaglutide on discrimination performance were specific to the alcohol discriminative stimulus, a new cohort of rats (7 males and 3 females) underwent discrimination training. This cohort was also used in Experiments 3 and 4. Given the limited number of females in this cohort, males and females are combined for Experiments 2-4. Upon testing, the lowest effective semaglutide dose from Experiment 1 (0.01 mg/kg) or vehicle was administered 3 h prior to water (IG). Testing was repeated such that all subjects received both semaglutide and vehicle, with at least two consecutive training days (including water and alcohol) with performance above 80%).

### Experiment 3: Tirzepatide

After completion of Experiment 2 and six intervening training days, rats began testing of tirzepatide (0, 0.1, 0.3 and 1.0 mg/kg, SC). Rats received the assigned dose of tirzepatide 3 h prior to alcohol (2 g/kg, IG) and then underwent a test session. Testing was repeated such that all subjects received each assigned dose, with a minimum of two training days with discrimination performance meeting the 80% criterion for each rat.

### Experiment 4: Retatrutide

After the completion of Experiment 3, rats returned to training for 18 sessions prior to testing with retatrutide. Testing procedures were identical to tirzepatide with the three doses of 0.03 mg/kg, 0.1 mg/kg, and 0.3 mg/kg.

### Data analysis

Statistical analyses were performed using GraphPad Prism (version 10.4), unless otherwise specified, with significance set at p < 0.05. The primary measure of discrimination accuracy was the percentage of alcohol-appropriate lever responding prior to delivery of the first sucrose reinforcer (i.e., the first 10 lever responses). Full expression of the interoceptive stimulus effects of alcohol was defined as ≥80% alcohol-appropriate responding. For Experiment 1, the alcohol cumulative curve and acute semaglutide injection data were analyzed using two-way ANOVAs, and repeated semaglutide injection data were examined via a three-way repeated-measures linear mixed model (in SPSS version 29.0.1.0). When sex emerged as a significant factor, separate ANOVAs were performed for males and females. In Experiment 2, due to limited sample size, data from both sexes were pooled and analyzed using one-way ANOVAs for cumulative curve and acute administration measures. Test days on which a rat failed to meet the FR10 criterion were excluded from primary analyses of %appropriate responding, but response rate data were retained to capture potential deficits in overall responding; therefore, when data were excluded mixed-effects models were used in these instances to incorporate all available data. Tukey’s HSD was employed for post hoc comparisons.

## Results

### Experiment 1

#### Cumulative curve

To confirm stimulus control an alcohol dose curve was determined prior to testing of semaglutide. A two-way repeated-measures ANOVA confirmed alcohol discriminative stimulus control, revealing a significant main effect of alcohol dose (F(3, 42) = 39.84, p < 0.0001), with both doses lower than the 2 g/kg training dose resulting in lower alcohol-appropriate responding (p < 0.01; Figure 1A). Response rate analysis also showed a significant main effect of alcohol dose (F(3, 42) = 3.68, p = 0.019), though post-hoc comparisons did not reveal significant differences among individual doses (Figure 1B). No sex differences were noted for either measure.

**Fig. 1:**
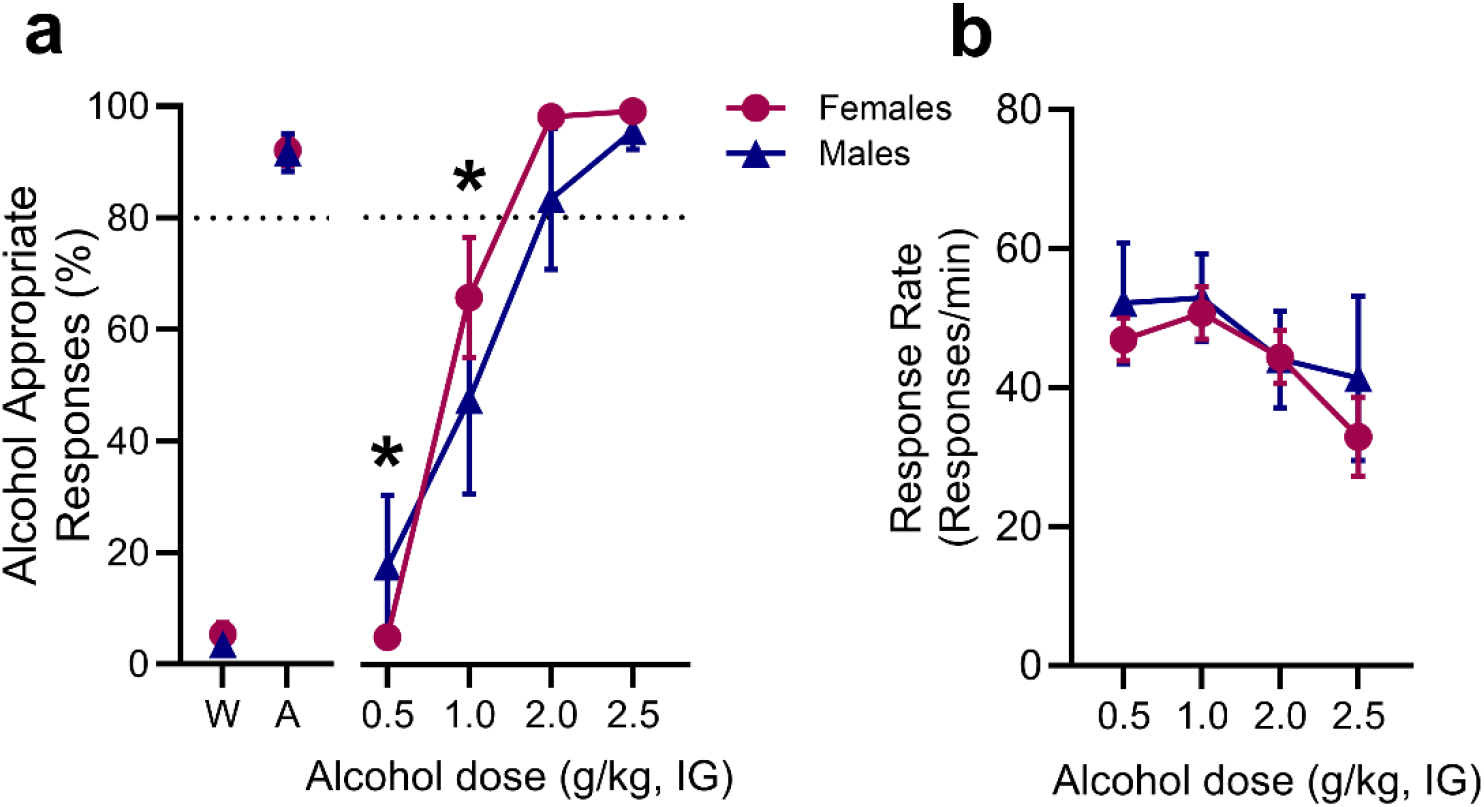
Prior to evaluating semaglutide, alcohol discriminative stimulus control was confirmed. (A) Average (±S.E.M.) alcohol-appropriate responses from the three alcohol and water sessions that preceded testing are represented to the left of the axis break. Substitution for the 2 g/kg alcohol training dose was observed, confirming stimulus control. Dotted line at 80% represents full substitution for alcohol training dose. (B) An overall reduction in response rate (responses/min) was observed. *significantly different from 2.0 g/kg training dose, p<0.05

#### Acute semaglutide

To evaluate the effects of semaglutide on discrimination performance, a 2-way repeated-measures ANOVA examining alcohol-appropriate responding revealed significant main effects of semaglutide dose (F(4, 48) = 15.98, p < 0.0001) and sex (F(1, 12) = 5.78, p = 0.02) as well as an interaction (F(4, 48) = 2.79, p < 0.05). As a main effect of sex was found, males and females were analyzed separately. In males, a main effect of semaglutide dose on alcohol-appropriate responding was found (F(4, 26) = 12.30, p < 0.0001), with the three highest doses reducing alcohol-appropriate responding (p<0.01; Figure 2A). Females similarly showed a significant main effect of dose (F(4, 22) = 6.81, p = 0.0006), with significant reductions at the two highest doses (p < 0.05; Figure 2B). A significant main effect of dose on response rate was observed in both males (F(4, 26) = 11.20, p < 0.0001) and females (F (4, 31) = 5.66, p < 0.01) and post-hoc analyses confirmed significant reductions in response rate at the highest dose tested compared to vehicle control in both sexes (p < 0.05; Figure 2C,D). These results show that acute semaglutide treatment effectively blunts the interoceptive effects of alcohol (2 g/kg).

**Fig. 2:**
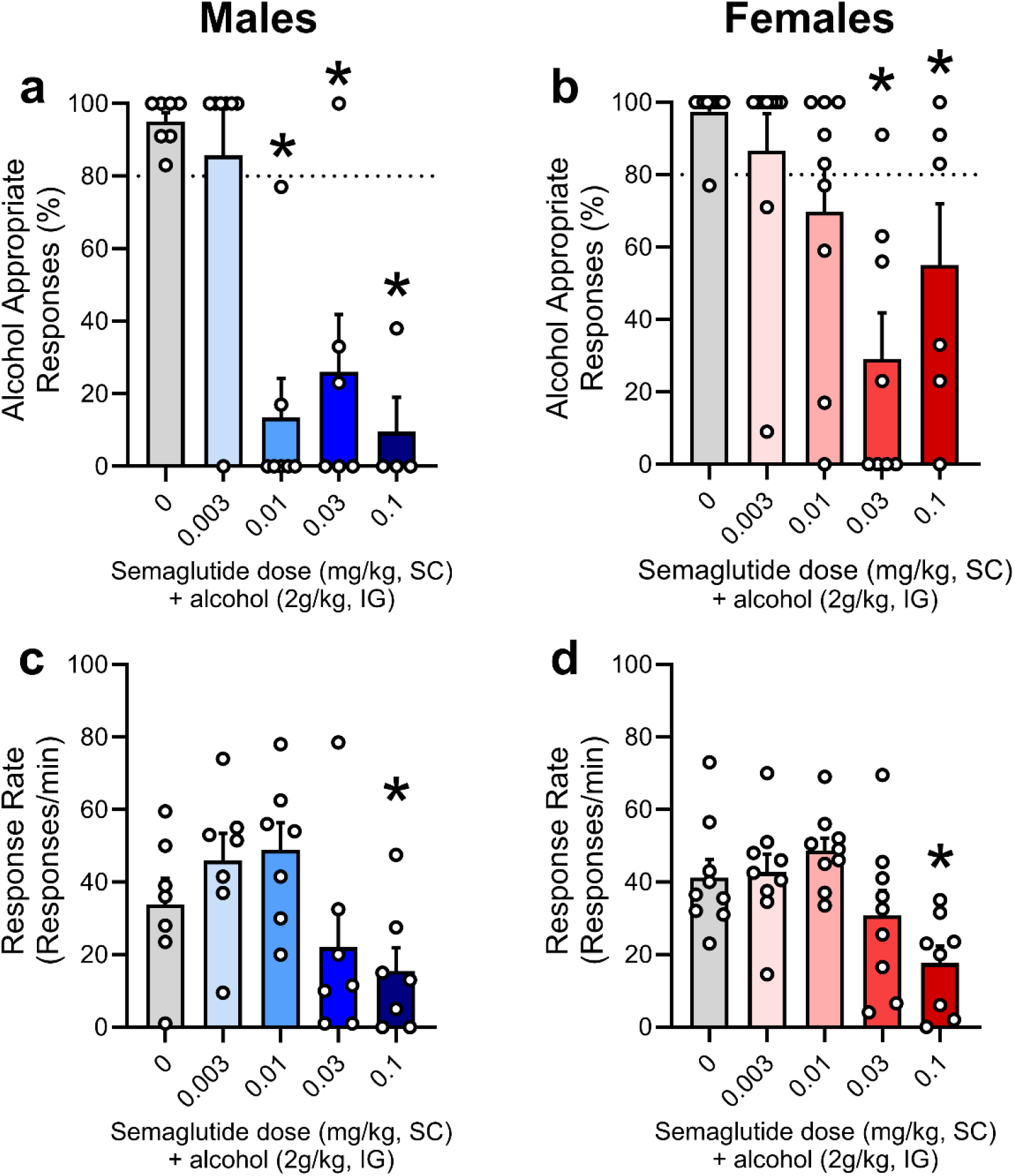
Experiment 1: Acute semaglutide. Semaglutide administered 3h prior to alcohol (2 g/kg) reduced alcohol-appropriate responses in both males (A) and females (B) in a dose-dependent manner. Dotted line at 80% represents full substitution for alcohol training dose. (C,D) Response rates were reduced in both sexes at the highest dose. *significantly different from vehicle, p<0.05

#### Repeated semaglutide

Next, we evaluated the effects of repeated semaglutide treatment on the discriminative stimulus effects of alcohol. A three-way repeated-measures linear mixed model revealed significant main effects of sex (F(1, 14.8) = 14.72, p <0.01), injection day (F(2, 140.5) = 3.84, p < .05), and dose (F(3, 141.2) = 36.01, p < 0.001), therefore data are shown as separate analyses for each sex. In both males and females, semaglutide treatment reduced sensitivity to the discriminative stimulus of alcohol. In males there was a significant main effect of semaglutide dose (F(3, 18) = 34.27, p < 0.0001) with significantly reduced alcohol-appropriate responding at all semaglutide doses relative to vehicle (ps < 0.0001). Post treatment testing found a main effect of injection day (F(1, 6) = 9.33, p < 0.05) but no effect of dose, and a significant injection day × dose interaction (F(3, 16) = 4.27, p < 0.05), where the highest 0.1 mg/kg dose reduced alcohol-appropriate responding only on the first post-injection day (Figure 3B). In females there was a main effect of dose (F(3, 24) = 9.45, p < 0.001) but no interaction, and post hocs indicated significantly reduced responding compared to vehicle at all doses except the lowest 0.01 dose (p<0.01). On post-injection days no effects were found in females (Figure 3C). Together this indicates that semaglutide altered interoceptive effects in both males and females with higher sensitivity in males, and these effects quickly returned to match vehicle when semaglutide was discontinued. A two-way ANOVA on male response rate during injections revealed significant main effects of injection day (F(2, 12) = 14.96, p < 0.001) and dose (F(3, 18) = 3.78, p < 0.05), as well as a significant injection day × dose interaction (F(6, 36) = 3.63, p < 0.01). Post-hoc comparisons showed that on injection day 1, only the highest dose (0.1 mg/kg) significantly reduced response rate compared to vehicle (p < 0.05; Figure 3D). On injection day 3, no dose significantly differed from vehicle. On injection day 8, the 0.01 mg/kg (p < 0.05) and 0.03 mg/kg (p < 0.05) doses differed from vehicle. No differences in response rate were found on post-injection days. In females there was a main effect of dose (F(3, 24) = 5.61, p < 0.01) and a significant injection day × dose interaction (F(6, 46) = 3.99, p < 0.01), where response rates were reduced at the highest dose (0.1 mg/kg) on injection day 1 (p < 0.01) and the lowest dose (0.01 mg/kg) differed on injection day 8. On post-treatment days there was a significant interaction (F(3, 24) = 3.82, p < 0.05) where the 0.03 dose had a higher response rate than vehicle on the first post treatment day (Figure 3E). Change in body weight by % was not significantly altered by semaglutide treatment in either males or females (Figure 3F, G), which may have been influenced by food restriction to 90%.

**Fig. 3:**
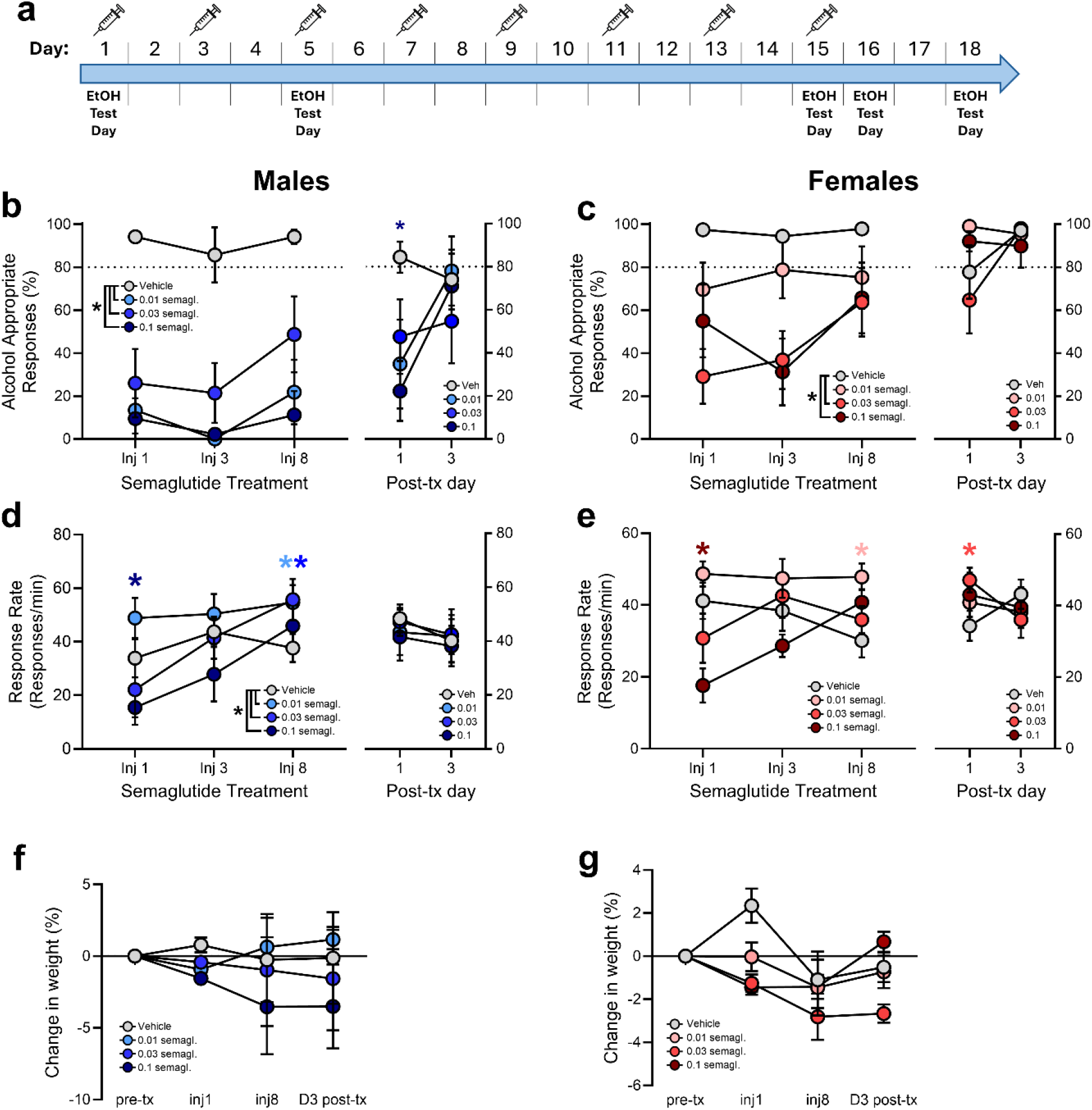
Experiment 1: Repeated semaglutide administered every-other-day for a total of eight injections, with alcohol discrimination tests occurring 3 h after the 1^st^, 3^rd^, and 8^th^ injection, as well as the 1st and 3rd post-treatment days (A). Semaglutide reduced alcohol-appropriate responses in both males (B) and females (C) in a dose-dependent manner. Dotted line at 80% represents full substitution for alcohol training dose. (D) Response rate was reduced in males after injection 1 at the highest dose. (E) Response rate was reduced in females at injections 1, 3, and 8, at the highest dose, and at injection 1 at the second highest dose. (F,G) Change in body weight (%) was not significantly altered by semaglutide treatment in males or females, respectively. *significantly different from vehicle, p<0.05

### Experiment 2

#### Cumulative curve

A one-way repeated-measures ANOVA confirmed stimulus control in cohort 2, revealing a significant effect of alcohol dose (F(3, 27) = 15.49, p < 0.0001), with both doses lower than the 2 g/kg training dose resulting in lower alcohol-appropriate responding (p < 0.01; Figure 4A). Alcohol dose had no effect on response rate (Figure 4B).

**Fig. 4:**
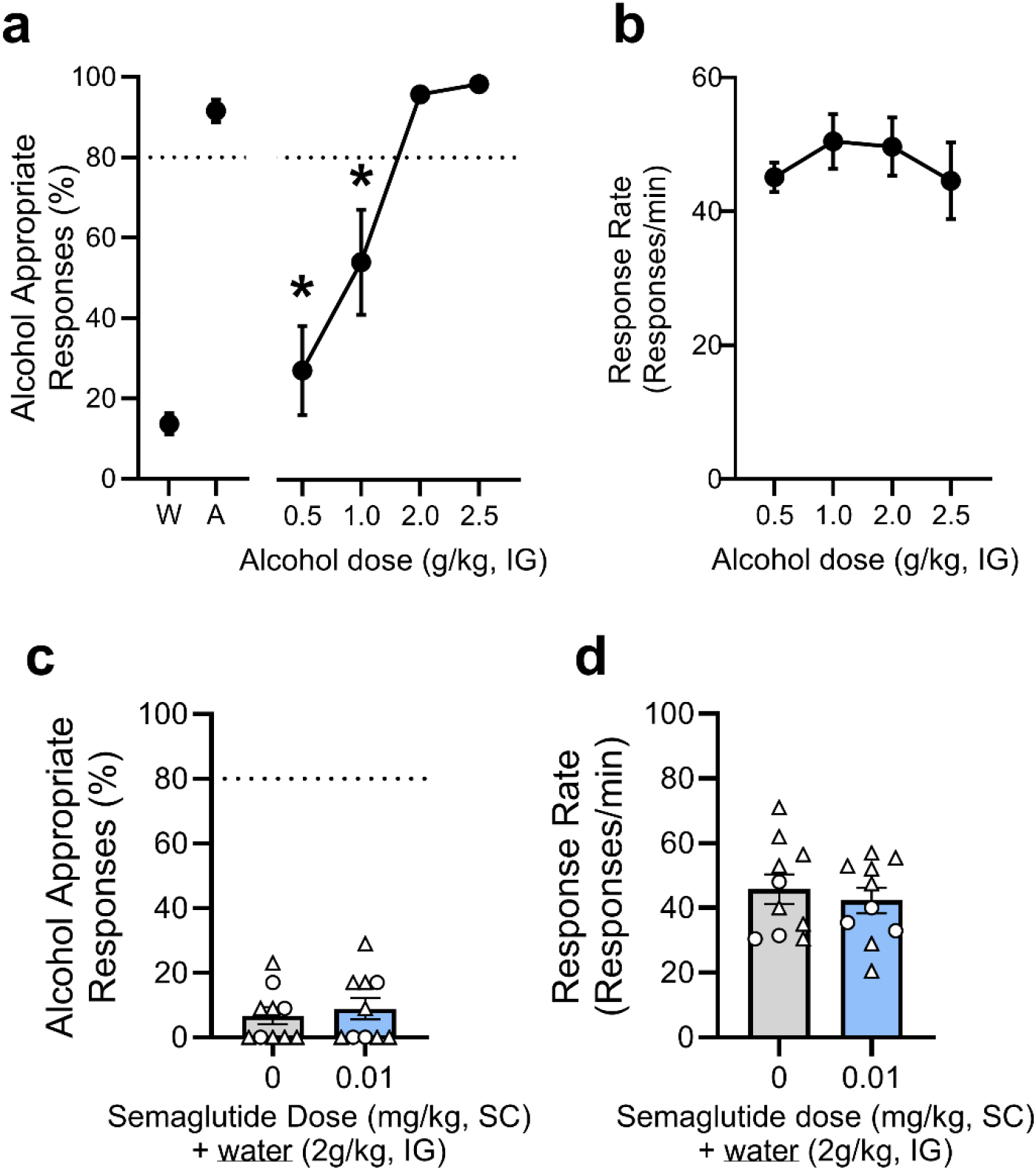
Experiment 2: Semaglutide on water sessions. (A) Prior to testing, alcohol stimulus control was confirmed as alcohol-appropriate responses were reduced at alcohol doses lower than 2 g/kg training dose, with no changes in response rate (B). Semaglutide administered 3 h prior to water had no effect on alcohol-appropriate responses (C) or response rate (D). Dotted line at 80% represents full substitution for alcohol training dose. ○ female subject, △ male subject. *significantly different from 2.0 g/kg training dose, p<0.05

#### Semaglutide on water session

Semaglutide had no effect on alcohol-appropriate responding or response rate on test days where subjects were administered water (Figure 4 C,D). This finding supports that semaglutide blunts the discriminative stimulus effect of alcohol without causing nonspecific changes in discrimination behavior.

### Experiment 3: Tirzepatide

Tirzepatide significantly reduced the discriminative stimulus effects of alcohol as confirmed by a one-way ANOVA on alcohol-appropriate responding that indicated a significant main effect of dose (F(3, 24) = 8.88, p < 0.001). Post-hoc analyses revealed significant reductions relative to vehicle at doses of 0.3 mg/kg (p < 0.05) and 1 mg/kg (p < 0.01; Figure 5A). On response rate there was a main effect of tirzepatide dose (F(3, 24) = 8.83, p < 0.001) with significantly reduced responding at only the highest dose (1 mg/kg, p < 0.01; Figure 5B). This indicates that the blunting of alcohol discriminative stimulus effects at the 0.3 mg/kg dose was not related to a nonspecific motor reduction.

**Fig. 5:**
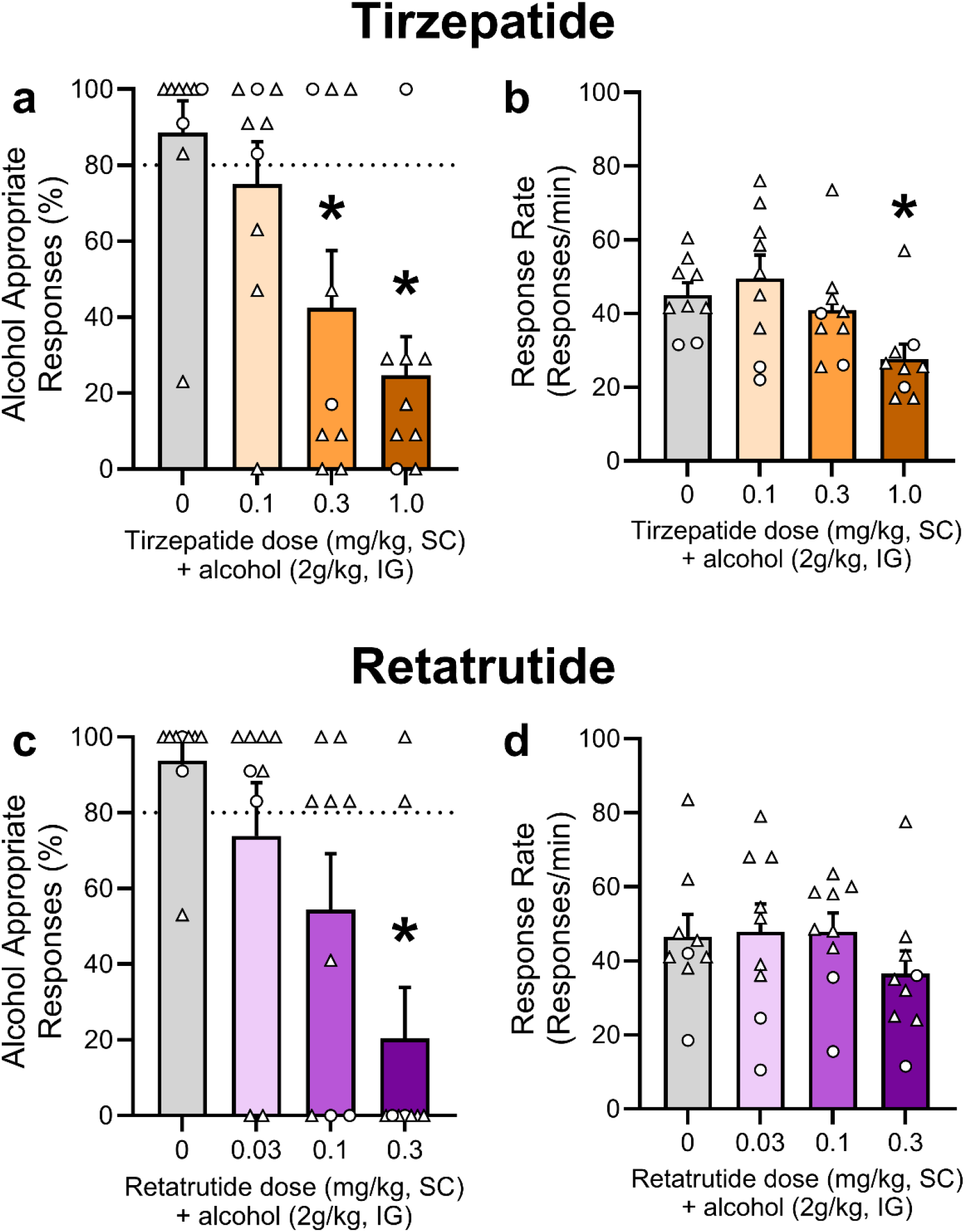
Experiment 3. (A) Tirzepatide administered 3 h prior to alcohol (2 g/kg) reduced alcohol-appropriate responding in a dose-dependent manner, and (B) only the highest dose reduced the response rate. (C) Retatrutide similarly reduced alcohol-appropriate responding while having no effect on response rate (D). Dotted line at 80% represents full substitution for alcohol training dose. ○ female subject, △ male subject. *significantly different from vehicle, p<0.05

### Experiment 4: Retatrutide

Retatrutide also significantly reduced the discriminative stimulus effects of alcohol as supported by a one-way ANOVA on alcohol-appropriate responding that found a significant main effect of retatrutide dose (F(3, 24) = 6.41, p < 0.01), with significant reduction in responding only at the highest dose tested (0.3 mg/kg) compared to vehicle (p < 0.01; Figure 5C). Retatrutide had no effects on response rate (Figure 5D).

## Discussion

Our findings demonstrate that GLP-1 receptor-based therapeutics significantly disrupt the interoceptive stimulus effects of alcohol, highlighting a potential therapeutic mechanism through which these medications may reduce alcohol consumption and motivation. Acute administration of semaglutide, tirzepatide, and retatrutide each dose-dependently attenuated alcohol discrimination, suggesting that blunting or altering subjective alcohol effects could be a common target of this drug class. By extending earlier preclinical work on GLP-1 receptor agonists and alcohol intake to a task that directly assesses the subjective effects of alcohol, these findings provide important context for interpreting clinical observations of reduced drinking behavior among individuals receiving GLP-1 receptor-based therapies.

In Experiment 1, acute semaglutide robustly disrupted alcohol discrimination, as evidenced by dose-dependent reductions in alcohol-appropriate responding in both sexes. The effective semaglutide dose range in this paradigm (0.01–0.1 mg/kg for males, 0.03-0.1 mg/kg for females) is consistent with doses reported to reduce alcohol self-administration and intake (0.0003-0.01mg/kg) in rodent and nonhuman primate models (0.003-0.01 mg/kg; Marty et al. 2020; Chuong et al. 2023; Aranäs et al. 2023; Fink-Jensen et al. 2024). Short-acting GLP-1 receptor agonists like exendin-4 and liraglutide have also been shown to disrupt conditioned place preference for alcohol supporting their ability to selectively attenuate the rewarding effects of alcohol (Egecioglu et al. 2013; Shirazi et al. 2013; Vallöf et al. 2016). In the present study, every-other-day treatment with semaglutide maintained efficacy across the 15-day period. However, by the final treatment day the strength of the effect had signs of waning at some doses, which suggested that if treatment were extended efficacy may drop off over time. This may reflect adaptive processes, consistent with clinical observations that initial effects, including adverse side effects such as nausea, often diminish with continued treatment (Wharton et al. 2022; Chao et al. 2023). This may explain why the highest dose of semaglutide reduced response rate only after the initial injection. A similar attenuation has been observed in long-term GLP-1 receptor agonist studies of alcohol intake in rodents, in which behavioral effects diminished across days after initial efficacy (Vallöf et al. 2016). Understanding whether similar adaptive processes influence therapeutic efficacy related to alcohol’s interoceptive effects will be an important area for further research. Importantly, semaglutide did not disrupt discrimination performance on water test days (Experiment 2), confirming that reductions in alcohol discrimination were specifically related to the attenuation of alcohol’s interoceptive cue rather than reflecting general impairment in performance. This specificity supports the interpretation that GLP-1 receptor agonists selectively target alcohol-related subjective effects.

Interestingly, the effectiveness of semaglutide differed between males and females, with males exhibiting greater sensitivity. The lowest effective dose in males (0.01 mg/kg) failed to significantly reduce alcohol-appropriate responding in females, suggesting lower sensitivity in females. In humans, women typically show greater responsiveness to GLP-1 receptor agonists compared to men, likely due to estrogen’s potentiation of GLP-1 receptor signaling within reward-related and metabolic pathways in the brain (Hayes et al. 2008; Finan et al. 2012; Richard et al. 2016; Rentzeperi et al. 2022; Badulescu et al. 2024). Notably, Richard et al. (2016) demonstrated that blocking estrogen receptor α signaling abolished the suppressive effects of GLP-1R activation on reward-related behavior, further supporting the idea that estrogen is a critical modulator of GLP-1’s central effects on motivated responding. Estrogen fluctuations differ markedly between species; female rats experience brief estrogen peaks during proestrus rather than the sustained elevations observed across the human menstrual cycle (Staley and Scharfman 2005; Richard et al. 2016). Thus, the transient nature of estrogen elevations in female rats may limit estrogen’s facilitatory effects on GLP-1 signaling, potentially explaining the more robust disruption of alcohol discrimination observed in males. This finding highlights the importance of considering sex differences and hormonal status in preclinical evaluations to enhance clinical translation.

In Experiments 3 and 4, we assessed acute treatment of the newer-generation GLP-1 receptor agonist therapeutics tirzepatide and retatrutide, both of which successfully attenuated alcohol discrimination. The broader receptor profile of these newer-generation GLP-1 receptor agonists is thought to offer clinical advantages, potentially providing better tolerability or enhanced effectiveness due to simultaneous modulation of reward-related neural circuits and metabolic pathways (Ray 2023; Tufvesson-Alm et al. 2023; Badulescu et al. 2024; Kaur and Misra 2024). In particular, retatrutide, a triple agonist targeting GLP-1 receptors, GIP, and glucagon receptors, has demonstrated substantial metabolic benefits, raising interest in its potential for treating substance use disorders as well (Jastreboff et al. 2023; Tufvesson-Alm et al. 2023; Tetelbaun et al.). These multi-target effects could potentially enhance therapeutic profiles compared to GLP-1 receptor agonists like semaglutide, and may differentially influence specific outcomes, such as alcohol intake and weight loss. Compounds that effectively reduce alcohol use while producing more modest effects in other domains may represent leading candidates for clinical development. While our data demonstrated clear efficacy for both compounds in reducing alcohol interoceptive effects across the tested dose ranges, it will be important in future work to assess efficacy following repeated treatment and to directly compare potential advantages in terms of efficacy, side-effect profiles, and tolerability. Furthermore, given the small sample sizes within sex for Experiments 3 and 4, sex comparisons could not be made. However, as we observed sex differences with semaglutide, assessing sex-specific responses to tirzepatide and retatrutide will be essential to fully characterize their therapeutic potential for AUD.

Drug discrimination procedures hold strong translational relevance, given their predictive validity for subjective drug effects in humans (Grant and Bennett 2003; Besheer and Hodge 2005; Solinas et al. 2006; Bolin et al. 2016; Lovelock et al. 2021). The subjective/interoceptive effects of alcohol significantly contribute to craving, consumption, and relapse, and thus represent an important therapeutic target (Grant and Bennett 2003; Besheer and Hodge 2005; Koob and Volkow 2010; Jaramillo et al. 2018b; Lovelock et al. 2021). By applying a reverse translational approach that evaluates compounds with emerging clinical efficacy, we demonstrated that semaglutide, tirzepatide, and retatrutide blunt alcohol interoceptive/subjective effects, potentially clarifying a clinically meaningful mechanism. These effects are consistent with user-reported reductions in alcohol effects and use on social media platforms (Arillotta et al. 2023, 2024; Quddos et al. 2023; Bremmer and Hendershot 2024). Future studies should directly assess whether these preclinical findings align with subjective-effect reductions in humans which could explain the reduced cravings reported during semaglutide treatment (Hendershot et al. 2025), further bridging animal and human data.

Collectively, these findings reinforce existing evidence that GLP-1 receptor agonists serve as promising candidates for treating AUD (Fink-Jensen and Vilsbøll 2016; Hayes and Schmidt 2016; Brunchmann et al. 2019; Klausen et al. 2022; Jerlhag 2023, 2025; Leggio et al. 2023; Bruns et al. 2024). By attenuating the subjective effects and reinforcing effects of alcohol, as supported by the present data and previous work, respectively (Egecioglu et al. 2013; Shirazi et al. 2013; Vallöf et al. 2016), these medications may help reduce alcohol motivation, intake, and relapse risk. As clinical use of GLP-1 receptor agonists continues to expand, understanding how these medications alter the subjective/interoceptive effects of alcohol will be critical for optimizing their utility in AUD. Human laboratory studies that incorporate subjective alcohol response assessments, alongside clinical outcomes, could provide a direct translational bridge (Koob et al. 2009) and clarify the mechanistic relevance of our findings.

